# Absence of reliable physiological signature of illusory body ownership revealed by fine-grained autonomic measurement during the rubber hand illusion

**DOI:** 10.1101/2020.07.24.219287

**Authors:** Hugo D Critchley, Vanessa Botan, Jamie Ward

**Affiliations:** School of Psychology, University of Sussex, UK; Sackler Centre for Consciousness Science, University of Sussex, UK; Brighton and Sussex Medical School, University of Sussex and University of Brighton UK

**Keywords:** Active inference, Autonomic, Illusion, Interoception, Selfhood

## Abstract

The neural representation of a ‘biological self’ is linked theoretically to the control of bodily physiology. In an infleuntial model, selfhood relates to internal agency and higher-order interoceptive representation, inferred from the predicted impact of efferent autonomic nerves on afferent viscerosensory feedback. Here we tested if an altered representation of physical self (illusory embodiment of an artificial hand) is accompanied by sustained shifts in autonomic activity.

Participants (N=37) underwent procedures for induction of the rubber hand illusion (synchronous stroking of own unseen hand and observed stroking of artificial hand) and a control condition (asychronous stroking). We recorded electrodermal activity, electrocardiography, and non-invasive measurement of multiunit skin sympathetic nerve activity (SKNA) from the chest. We compared measures between task conditions, and between individuals who did and did not experience the illusion. Bayes factors quantified the strength of evidence for and against null hypotheses.

**S**ubjective reports and observed proprioceptive drift confirmed the efficacy of synchronous vs asynchronous condition in inducing illusory hand ownership. Stringent discriminant analysis classified 24 /37 individuals as experiencing the rubber hand illusion. Electrodermal activity, heart rate, heart rate variability, and SKNA measures revealed no autonomic differences between synchronous vs asynchronous conditions, nor between individuals who did or did not experience the rubber hand illusion. Bayes factors confirmed there was substantial evidence for no physiological differences.

In contrast to earlier reports, our cardiac and electrodermal autonomic data, including fine grained SKNA measurement, provide substantial evidence for the absence of a reliable change in physiological state during induction or experience of the rubber hand illusion. If illusory body ownership is coupled to, or facilitated by, changes in efferent autonomic activity and afferent viscerosensory feedback, our findings suggest that such changes in bodily physiology are not sustained as an obligatory component of the illusion.

## Introduction

The representation of the body’s physiological state and physical boundaries is argued to be fundamental to self-perception and awareness. This representation is built upon the integration of sensory information across modalities with expectations and predictions regarding what our body should be doing [1–4]. The dependence of the experience of body ownership on dynamic coherence across senses is illustrated by the rubber hand illusion, induced through correspondence between somatosensory stimulation and visual feedback: An artificial ‘rubber’ hand is placed in front of a participant and seen to be stroked. At the same time, the matching own hand of the participant, is hidden from sight, yet stroked at the same frequency (synchronously). Most participants experience the rubber hand as part of their own body [4]. This experience is reported by the participant as a subjective feeling, the strength of which can be scored on questionnaires and visual analogue scales. Less subjective measures of the illusion include ‘proprioceptive drift’, where the judged location of the participant’s own hand shifts to be nearer the rubber hand. Physiological responses can also provide objective measurement of the rubber hand illusion: for example when the rubber hand is threatened by, for instance being hit with a hammer, the strength of illusory embodiment can be inferred by the magnitude of an autonomic reaction to the threat, such as skin conductance response [5] or motoric withdrawal of the participant’s own hand [6].

Influential theories of consciousness suggest that there is a primacy to the (predictive) control of internal bodily state, through which the sense of self as a continuous, unitary, bounded experience arises from the inseparability of mental processes from the dynamics of interoceptive (inner physiological; viscerosensory) sensation and internal agency [7–9]. Empirical data to support this is limited. Abnormalities in normative autonomic reactivity or interoceptive functioning are associated with clinical conditions involving disturbed self-representation [10,11]. Furthermore, individuals with reduced ability to perform a heartbeat tracking task (a heuristic measure of sensitivity to interoceptive signals) are more susceptible to the rubber hand illusion [12], suggesting that a weaker internally driven model of self predisposes to a more malleable representation of the body’s boundaries. Relatedly, if physiological information is introduced into the display a virtual rubber hand ([13] or body [14] by colour changes pulsing normatively in time with the participant’s own heartbeat, this can increase the likelihood (and objective correlates of) the illusory experience of body ownership, indicating a binding role of interoceptive predictive experience in the representation of the physical self.

Within active inference models of interoception, autonomic drive to peripheral organs encompasses predictions about the desired internal state, changing the afferent feedback to better manage interoceptive prediction error. As noted, interoceptive predictions and internal agency are central to theoretical models of embodied selfhood. By extension, one can hypothesize that changes in peripheral physiological state, mediated autonomically, would reflect a shift in bodily self-representation associated with the experience of the rubber hand illusion. Such changes are distinct from measuring reactions to threat provocation, in that they are more proximate to the putative interoceptive mechanism underpinning conscious selfhood. Relevantly, a fall in temperature of substituted (participant’s own stroked) hand is described as an objective indicator of the illusion of embodiment ([15]. Conversely, cooling the tested hand may facilitate the experience of the illusion [16]. In a stable environment, cooling of the skin surface can occur through changes in skin blood flow (perfusion) and increased sweat production through sweat gland activity (related to electrodermal activity measures such as skin sympathetic nerve activity). Immobility may be a likely trigger: Such local autonomic changes therefore can be viewed as an incidental passive consequence of a state of relative proprioceptive quiescence, amplified by the suggested / intentional state evoked by the active (versus asynchronous stroking) rubber hand illusion induction procedure. Thus, the physiological state of the participant’s own hand better matches the cold lifeless visual appearance of the embodied rubber hand. This finding of skin cooling during the rubber hand illusion has not been reliably replicated [17, 18]. Recently, however electrodermal activity (sympathetic skin responses) is reported to be sensitive rubber hand illusion, with variance of skin conductance differentiating between synchronous stroking condition (when illusion occurs) versus asynchronous control condition, in a rather transient way suggesting that the novelty of the experience is a potential psychological cause [19].

Electrodermal activity / skin conductance response is a measurable expression of skin sympathetic activity regulating eccrine sweat gland function, through both phasic responses and tonic shifts [20, 21]. Electrodermal activity is usually measured on the palmar skin of the hand and is a sensitive and widely used index of central attention and arousal states. Nevertheless (of relevance to interoception), skin sweat glands do not possess dedicated visceral afferents, hence sensory feedback of electrodermal activity is absent or indirect (via correlated changes in physiological arousal in other organ systems; but see [22]). Organ specificity also exists: The autonomic sympathetic innervation of the palmar skin is distinct from the sympathetic innervation of vasculature and heart [23]. The heart (and its atrioventricular pacemaker) also has an antagonistic autonomic input via cardiovagal parasympathetic fibres and is, unlike electrodermal activity, is subject to baroreflex control: The rate and strength of cardiac contraction via autonomic efferents are determined by afferent feedback from arterial baroreceptors in the aorta and carotids. Heart rate and heart rate variability thus provide indices of dynamic interaction between such autonomic efferent activity and interoceptive cardiac feedback.

Here, participants underwent a rubber hand illusion induction protocol, involving synchronous stroking of a visible artificial right hand and the participants own hidden right hand, and a control comparison condition in which stroking of the two hands was asynchronous. To quantify changes in autonomic reactivity that we predicted would reflect changes in body ownership (experience of the rubber hand illusion), we recorded; 1) electrodermal activity from the participants right hand (to measure magnitude and frequency of sympathetic skin responses); 2) electrocardiography (to measure heart rate and vagally-mediated heart rate variability, HRV), and lastly; 3) a relatively novel non-invasive purported measure of skin sympathetic nerve activity (SKNA) recorded over the chest in a distribution reflecting sympathetic activity of the stellate ganglion [24–26]. This SKNA measure was included as a fine-grained direct measure of efferent autonomic nerve traffic. We tested the hypothesis that cardiac, electrodermal, and SKNA autonomic indices would differentiate the synchronous induction vs asynchronous control conditions in the rubber hand illusion protocol, and between individual who experienced the illusion to a greater extent. We further tested for association with subjective and objective measures of the illusion and with individual differences in interoceptive sensitivity and awareness indexed by heartbeat counting performance and metacognitive insight.

## Materials and Methods

### Participants

A total of 37 participants (mean age=21.59, SD=3.69; 31 females) completed the study. Participants were recruited from students and staff at the University of Sussex via electronic advertisement. Ethical approval was obtained from the Brighton and Sussex Medical School Research Governance and Ethics Committee (BSMSREGC) and the Science and Technology Research Ethics Committee of the University of Sussex. All participants give written informed consent.

### Rubber hand illusion paradigm

Testing was conducted in a climate-controlled room. In the rubber hand illusion task, the participant’s right arm was placed in a box (86 cm×60 cm×20 cm), screened from view. A life-size model of a right hand was placed midline within the visible section of the box, directly in front of participant body. The distance between the participant’s right index finger and the index finger of the fake hand was 20 cm.

Stroking with a paintbrush was applied by the experimenter (VB) to the index finger of the participants hidden hand and to the artificial hand. Two conditions were performed in a counterbalanced order across participants: 1) Synchronous stroking when the timing of the brush strokes on the participant’s own hand and on the rubber hand was coincided. 2) Asynchronous stroking when the timing of the brush strokes on the participant’s hand and rubber hand was out of phase by approximately 625ms). Each condition lasted two minutes. At the beginning of each condition, the participant estimated the location of her/his right index fingertip three times by reading the corresponding number along a one-meter ruler, whose visible position varied each time to prevent the participant repeating responses in subsequent readings. Post-induction finger location judgements were obtained in the same manner as prior to the induction. Proprioceptive drift was calculated by subtracting the average of the pre-induction finger location judgements from the average of post-induction finger location judgement:

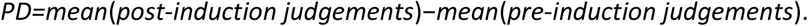

After each condition, the participant completed the RHI questionnaire comprising 10 items divided into three subscales: ownership, location, and agency [27], see *Table 1* for further details. The items were measured on a 7-point Likert scale (1=strongly disagree, 7=strongly agree) and the average score for each subscale was calculated.

**Table 1.**
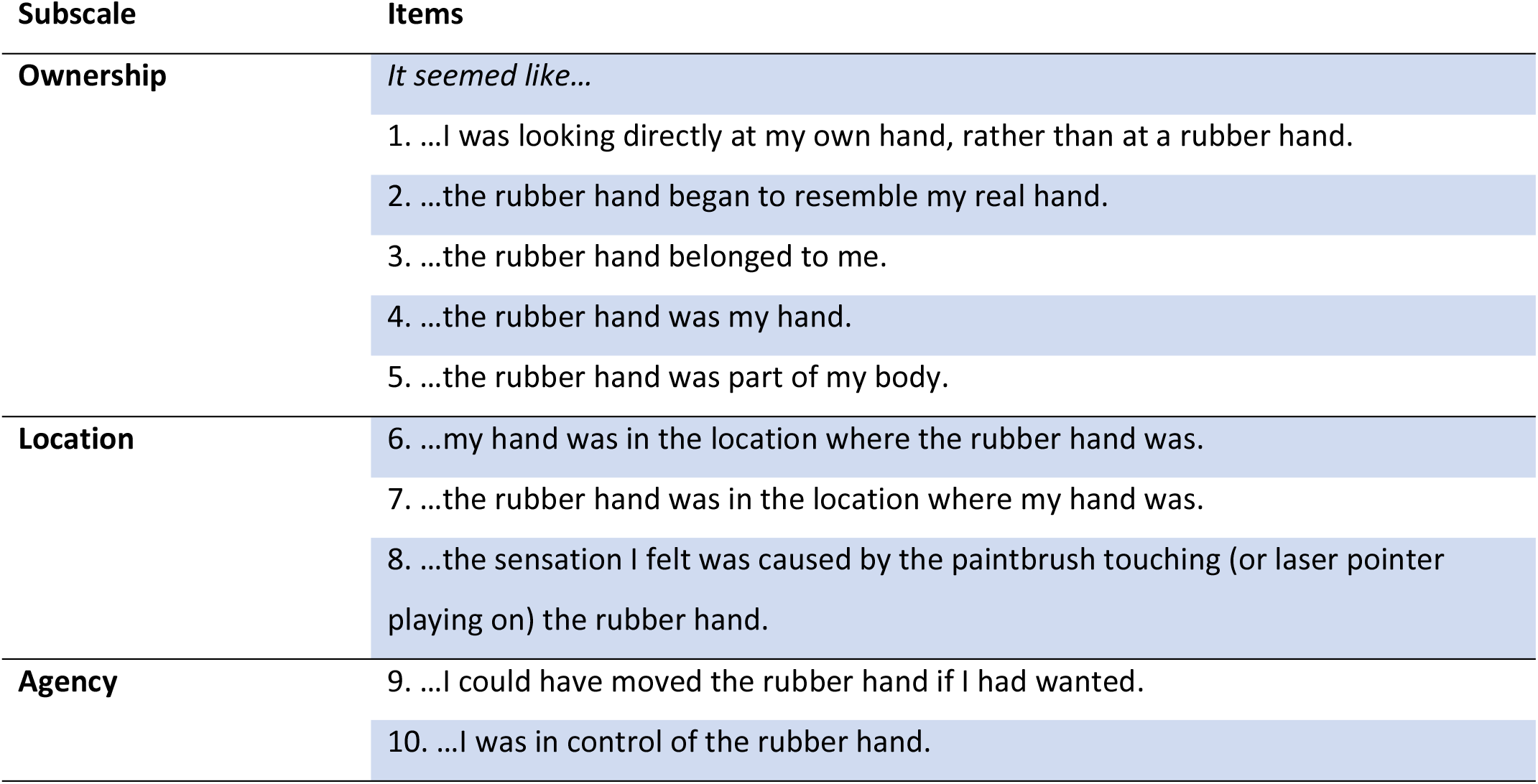
Rubber hand illusion questionnaire items and subscales. (See [27])

#### Interoceptive accuracy and awareness

Interoceptive accuracy was measured using the heartbeat tracking task [28] containing six trials with varying interval durations of 25, 30, 35, 40, 45 and 50 seconds played in a randomised order. The participant was instructed to silently count the number of heartbeats perceived in the given interval and to report them at the end of each trial. The actual number of heartbeats was measured using medical grade pulse oximeter with a soft sensor (Nonin Medical Inc, Plymouth MN USA; Xpod 8000S). For each trial, the accuracy score was derived using the following formula:

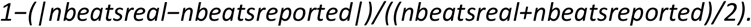

The resulting scores of each trial were averaged yielding the overall value for each participant [29]. Confidence judgements were taken at the end of each trial, participants being asked to rate the confidence they had in their reported number of heartbeats. Their response was recorded on a 10 points continuous visual analogue scale (VAS) from ‘total guess/no heartbeat awareness’ to ‘complete confidence/full perception of heartbeat’. Interoceptive awareness was then calculated using the Pearson correlation between interoceptive accuracy and confidence rating [29].

### Physiological measures

#### Heart rate and heart rate variability

Electrocardiographic signals were recorded using CED 1902 and 1401 hardware and fed into Spike 2 software (Cambridge Electronic Design Ltd; Cambridge UK) applying a 10Hz high bandpass filter and 100 Hz low bandpass filter [30]. For the analysis, a threshold was applied to isolate R-wave peaks and to extract the number of heartbeats in a given time interval. This gave measures of heart rate (HR) (beats per time interval) and heart rate variability (HRV) expressed as the root mean square of successive differences (RMSSD) between normal heartbeats, the primary time-domain measure for short-term variation, strongly correlated with high-frequency variations and an indicator of the vagally mediated (parasympathetic) changes reflected in HRV [31, 32]. Both HR and HRV were calculated for synchronous and asynchronous conditions.

#### Electrodermal activity (skin conductance responses)

Electrodermal activity was recorded using two finger electrodes (CED2502) and electrolyte gel placed on the palmar surface of the index and middle finger of the participant’s right hand. Signals were recorded into Spike 2 software using CED 1401 hardware. Analyses of skin conductance responses (SCRs) were performed on data exported to Matlab using Ledalab (V3.4.9) software. Adaptive data smoothing was applied, and continuous decomposition analysis was performed with extraction of continuous phasic and tonic activity. All SCR-onsets and amplitudes of above-threshold SCRs (a minimum of 0.01 microS) were then used to compute the average SCR amplitude per condition and a total number of SCR peaks per condition.

#### Skin sympathetic nerve activity (SKNA)

Following methods described by Doytchinova and colleagues [24], six electrocardiographic electrodes were placed over the chest of the participant, two placed on the wrists, and two placed on the ankles (Figure 1). This signal was captured with PowerLab 16/35 (ADInstruments, Dunedin NZ) and recorded and displayed using LabChart v7.

**Figure 1.**
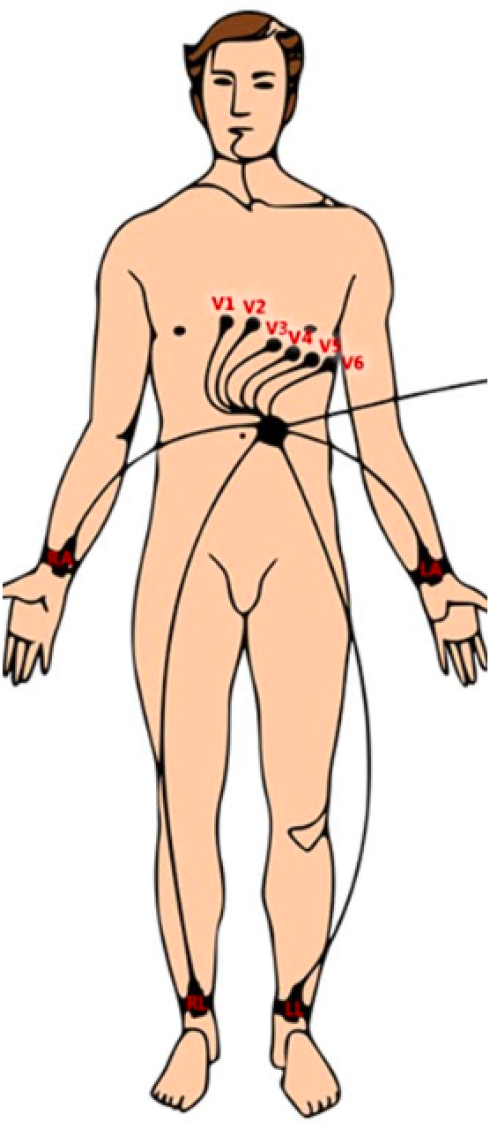
Electrode positioning for recording of skin sympathetic nerve activity (SKNA). To record SKNA, we followed methods described by Doytchinova and colleagues [24]: Electocardiographic (ECG) patch electrodes were used to record high frequency electrical signals banpass filtered between 500Hz and 1000Hz and derive the average signal (aSKNA) for each of the two experimental conditions. Correspondance between aSKNA activity and stellate ganglion function has been established in animal and human studies (reviewed [26]).

The sampling rate was set at 10000 samples/s and bandpass filtering between 500Hz and 1000Hz was applied. This filtering is shown to provide the best signal-to-noise ratio for skin sympathetic nerve activity (SKNA) recordings [24,26]. Quantitative signal data analysis was conducted, following the published protocol [24]. Here, the average SKNA (aSKNA) was calculated for specific time windows by dividing the total voltage within a time window to the number of samples within that specific time window. We used 30s-time windows and a sampling rate of 10,000/s. The total number of samples within the time window was 300,000. The total voltage of all the samples in the time window was then divided to 300,000. The aSKNA was then used in the statistical analyses presented in the Results section below.

#### Statistical analyses

Final statistical analyses were run in SPSS25 (IBM Statistics). Independent samples t-tests and their Bayesian equivalent (i.e. independent samples normal) tests were run on proprioceptive drift and subjective ratings for two groups (low vs high interoceptive accuracy) of seventeen participants each. Paired sample t-tests and their Bayesian equivalent (i.e. related samples normal) tests were performed to compare physiological measures between conditions. Pearson correlations and their Bayesian equivalents were also run to establish the relation between drift and interoceptive accuracy and awareness. Equal variances were assumed and Rouder’s method was applied for the Bayesian analyses. When normality was not assumed (i.e. questionnaire data and aSKNA data), non-parametric tests were run to re-confirm the results (see Supplementary material).

A *discriminant analysis* was run to distinguish between participants who did and did not show a strong experience of the illusion, using the difference between synchronous and asynchronous conditions in subjective ratings and in proprioceptive drift as predictors. The correlations between these predictors was relatively low (r=0.367) and Wilks’ Lambda was statistically significant for both predicting variables (p<0.001), confirming their adequacy for the analysis. Following this discriminant analysis, independent samples t-tests and their Bayesian equivalent (i.e. independent samples normal) tests were used to compare aSKNA results between the two groups of participants classified according to strength of their experience of the illusion.

## Results

### Behavioural results

#### Proprioceptive drift

Proprioceptive drift data was normally distributed for both synchronous and asynchronous drift measures (Shapiro Wilk normality tests; p=0.091 and p=0.945 respectively) and normality Q-Q plots. For the entire sample, proprioceptive drift was greater in the synchronous condition (M=26.0 mm, SD=5.46) than in the asynchronous condition (M=4.6mm, SD=3.67), t(36)=3.097, p=0.004, 95% CI [7.38, 35.41], d=7.56, B_H(0,1)_=0.123, thus there is moderate evidence for H1. Therefore, according to a typical control procedure, the RHI was successfully induced (Figure 2).

**Figure 2.**
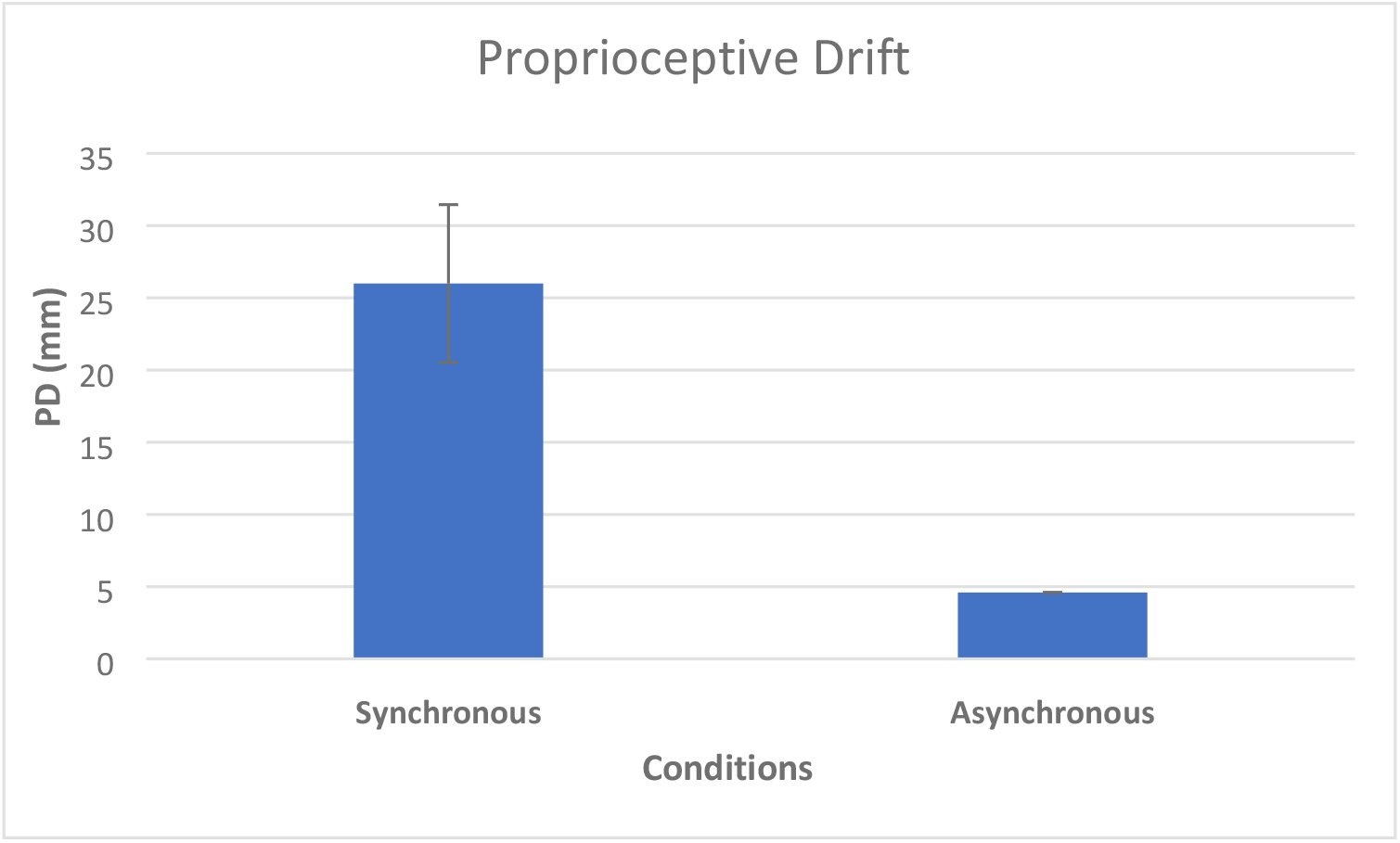
Proprioceptive drift results for synchronous and asynchronous conditions expressed as mean ± 1SE. Each participant judged the location of their unseen index finger on prior to and after each synchronous and asynchronous stroking condition (synchrony relative to the observed stroking of a visible artificial ‘rubber’ hand placed midline, directly in front of the participant). Proprioceptive drift is the objective finding that the experience of the illusion is accompanied by the participant judging his/her own hand to be located nearer to the artificial hand. The amount of measure ‘drift’ in a participant’s rating of his/her unseen hand is thus used as an objective measure of the illusion. Here, this measure of embodiment shows the synchronous condition induced the illusory experience.

### Subjective reports

The three subscales of the RHI questionnaire were independently analysed providing the following results: ownership, t(36)=7.825, p<0.001, 95% CI [1.23, 2.10], d=1.28, B_H(0,1)_=1.0 × 10^−3^; location, t(36)=6.187, p<0.001, 95% CI [0.87, 1.72], d=0.73, B_H(0,1)_=1.0 × 10^−3^; agency, t(36)=5.036 p<0.001, 95% CI [0.87, 2.00], d=0.85, B_H(0,1)_=1.0 × 10^−3^. As such, there was decisive evidence for H_1_, indicating that the illusion was successfully induced. (Table 2).

**Table 2.** showing means and standard deviations for subjective ratings in each condition and for each subscale.

|  | Synchronous | Asynchronous |
| --- | --- | --- |
| Ownership | $4.31 \pm 1.95$ | $2.65 \pm 1.61$ |
| Location | $4.14 \pm 1.46$ | $2.85 \pm 1.33$ |
| Agency | $3.53 \pm 1.87$ | $2.11 \pm 1.38$ |

#### Discriminant analysis

Participants with greater experience of the illusion could be distinguished by discriminant analysis from those with weaker or no experience of the illusion. There was a medium correlation between predictors (proprioceptive drift and subjective ratings) (r=0.367) and Wilks’ Lambda was statistically significant for both predicting variables (p<0.001), confirming their adequacy for the analysis. An Eigenvalue (=2.220) explained 100% of the variance with a high canonical correlation of 0.830. The model indicated two groups (illusion vs non-illusion) and had a high sensitivity (91.7%) and specificity (100%). The discriminant analysis indicated that twenty-four participants experienced the illusion (21 Females, M=22.15 yrs, SD=4.81) and thirteen participants had minimal experience of the illusion (10 Females, M=21.29 yrs, SD=3.00). The groups did not differ by age (t(35)=0.763, p=0.505), nor gender (χ^2^=0.694, p=0.405).

### Interoceptive accuracy

To test whether individual differences in interoceptive sensitivity influenced the experience of the rubber hand illusions in this sample [12], first a median split was used to divide participants into high interoceptive accuracy and low interoceptive accuracy groups [33]. We then tested for group differences between in strength of the rubber hand illusion. During synchronous proprioceptive drift, there was substantial evidence for no observable difference between high and low interceptive accuracy groups (t(32)=−0.734, p=0.468, 95% CI [−15.48, 32.94], d=0.25, B_H(0,1)_=3.183). Similarly, there was substantial evidence for no difference for asynchronous proprioceptive drift, arguably a control for ‘suggestibility’ (Longo et al., 2008); (t(32)=0.466, p=0.644, 95% CI [−12.56, 20.02], d=0.16, B_H(0,1)_=3.656)

Regarding subjective ratings in the synchronous condition, there were no differences on any of the subscales: ownership (t(32)=1.132, p=0.266, B_H(0,1)_=2.323); location (t(32)=−0.344, p=0.733, B_H(0,1)_=3.816); agency (t(32)=−0.225, p=0.824, B_H(0,1)_=3.929). Regarding subjective ratings in the asynchronous condition, there were no differences on any of the subscales: ownership (t(32)=0.983, p=0.334, B_H(0,1)_=2.252); location (t(32)=−0.249, p=0.805, B_H(0,1)_=3.910); agency (t(32)=−0.239, p=0.813, B_H(0,1)_=3.918). Again, for both synchronous and asynchronous conditions, Bayes factors showed the evidence for no interoceptive group effects to be substantial for location and agency while the measures for ownership was anecdotal [34].

Correlation analyses were run between drift measures and interoceptive accuracy and awareness scores. There was substantial evidence for no correlation between interoceptive accuracy and synchronous drift (r=−0.277, p=0.197, B_H(0, 1)_=3.284), asynchronous drift (r=−0.138, p=0.436, B_H(0, 1)_=5.561), nor the difference between synchronous and asynchronous drift (r=−0.107, p=0.547, B_H(0, 1)_=6.273). The same trend was observed for interoceptive awareness and synchronous drift (r=−0.064, p=0.721, B_H(0, 1)_=7.050), asynchronous drift (r=−0.097, p=0.585, B_H(0, 1)_=6.479), and the difference between synchronous and asynchronous drift (r=−0.001, p=0.996, B_H(0, 1)_=7.512). Together, these results indicate no significant difference between low- and high-interoceptive perceivers in the strength of the illusion as indicated by proprioceptive drift and/or subjective ratings.

### Physiological results

#### Heart rate and heart rate variability differences between synchronous and asynchronous conditions

Across the entire population, heart rate during the synchronous rubber hand induction condition was shown (substantial evidence) to be equivalent to that observed in the asynchronous condition (t(36)=0.088, p=0.930, 95% CI [−1.84, 2.001], d=0.299, B_H(0,1)_=7.79). Heart rate variability (RMSSD) during synchronous and asynchronous conditions was also similar (anecdotal evidence; t(36)=1.425, p=0.163, 95% CI [−2.41, 0.95], d=0.221, B_H(0,1)_=2.97). Together, these results indicate that cardiac physiology ‘sympathetic’ heart rate and parasympathetic RMSSD did not differentiate the synchronous stroking condition, associated with the induction and experience of the rubber hand illusion from asynchronous ‘illusion free’ control condition.

#### Heart rate and heart rate variability differences between participants who got the illusion and those who did not

For the synchronous condition, we observed no differences in heart rate between participants who did and did not experience the rubber hand illusion (anecdotal evidence; t(34)=−1.423, p=0.164, 95% CI [−12.86, 1.79], d=5.18, B_H(0,1)_=1.70]. The same was true for heart rate variability (substantial evidence; t(34)=−0.195, p=0.846, 95% CI [−21.2, 17.7], d=0.067, B_H(0,1)_=3.94). For the asynchronous condition, there were also no group differences in heart rate (anecdotal evidence; t(34)=−1.344, p=0.188, 95% CI [−13.6, 0.255], d=0.48, B_H(0,1)_=1.80) nor heart rate variability (substantial evidence; t(34)=0.786, p=0.437, 95% CI [−9.88, 21.7], d=0.28, B_H(0,1)_=3.07). Together, these results indicate with moderate evidence that cardiac physiology ‘sympathetic’ heart rate and parasympathetic RMSSD did not differentiate between individuals who did and did not experience the rubber hand illusion.

#### Electrodermal activity differences between synchronous and asynchronous conditions

We observed no significant difference between the synchronous and asynchronous conditions in the mean amplitude of skin conductance responses (substantial evidence; t(36)=−0.988, p=0.330, 95% CI [−2.45, 12.92], d=0.206, B_H(0,1)_=4.88); nor frequency (substantial evidence; t(36)=−0.911, p=0.369, 95% CI [−2.45, 12.92], d=0.089, B_H(0,1)_=5.235).

#### Electrodermal activity differences between participants who experienced the illusion and those who did not

For the synchronous condition, we observed no differences in mean amplitude of skin conductance responses (SCRs) between participants who did and did not experience the rubber hand illusion (very strong evidence; t(34)=−0.561, p=0.578, 95% CI [−0.76, 0.051], d=3.36, B_H(0,1)_=13.50). The same was true for frequency of SCRs (substantial evidence; t(34)=−0.373, p=0.711, 95% CI [−6.51 14.60], d=0.13, B_H(0,1)_=3.94). For the asynchronous condition, there were also no observed group differences in mean amplitude of SCRs (substantial evidence; t(34)=0.60, p=0.550, 95% CI [−0.047, 0.094], d=0.22, B_H(0,1)_=3.42), nor number of SCRs (substantial evidence t(34)=−0.21, p=0.983, 95% CI [−6.17, 6.03], d=0.01, B_H(0,1)_=4.00). Together, these results indicate that sympathetic electrodermal activity (SCRs) did not differentiate the synchronous stroking condition, associated with the induction and experience of the rubber hand illusion from asynchronous ‘illusion free’ control condition. Moreover, SCRs did not differ between those individuals who experienced the illusion from those that did not.

#### SKNA differences between synchronous and asynchronous conditions

The aSKNA corresponding to a time-window of 30s seconds was used to investigate differences between synchronous and asynchronous conditions. There were four time-windows for each condition (30s, 60s, 90s, and 120s) which were compared against each other. The results obtained are as follows: 30s, t(36)=−1.311, p=0.198, 95% CI [−0.02, 0.004], d=0.011, B_H(0,1)_=3.441; 60s, t(36)=−0.741, p=0.464, 95% CI [−0.02, 0.009], d=−0.07, B_H(0,1)_=5.991; 90s, t(36)=−0.674, p=0.505, 95% CI [−0.01, 0.007], d=0.05, B_H(0,1)_=6.272; 120s, t(36)=0.053, p=0.958, 95% CI [−0.01, 0.01], d=0.01, B_H(0,1)_=7.809. Therefore, there is strong evidence for H_0_, indicating that there are no differences in aSKNA between the two conditions (Figure 3).

**Figure 3.**
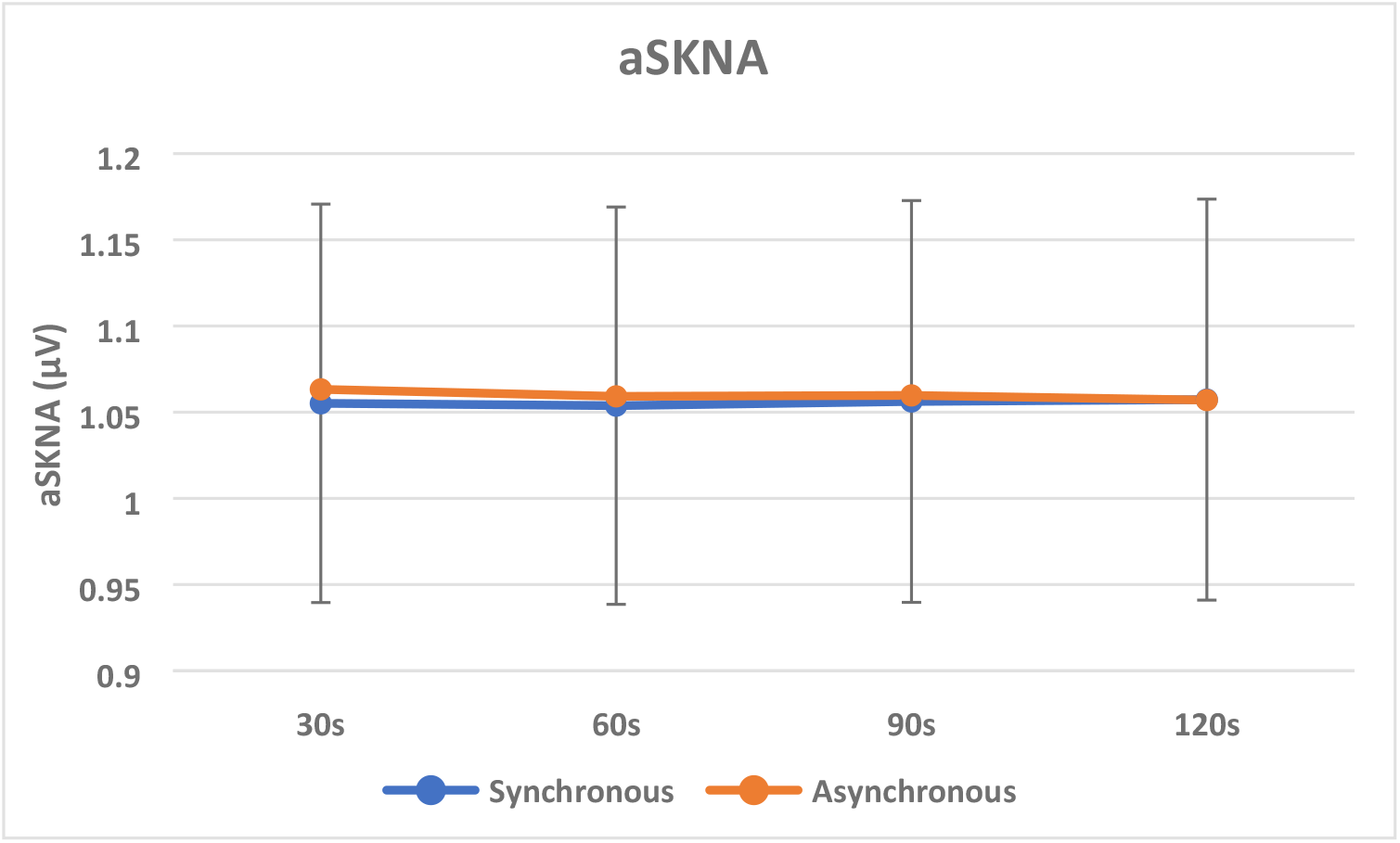
Average SKNA in synchronous and asynchronous conditions for each 30 s time-window expressed as means ± 1SE. Following the methods of Doytchinova and colleagues [24], continuous recording of SNKA was processed to give average scores over the 30s for each synchronous (active – associated with rubber hand illusion) and asynchronous (control – not associated with the rubber hand illusion) stroking conditions of the task. No consistent differences between conditions were observed across participants.

#### SKNA differences between participants who got the illusion and those who did not

The aSKNA differences between participants who got the illusion and those who did not as indicated by the discriminant factor analysis were analysed for each condition and each time window. For the synchronous condition, the results are as follows: 30s, t(35)=−0.366, p=0.716, 95% CI [−0.59, 0.40], d=0.13, B_H(0,1)_=3.777; 60s, t(35)=−0.368, p=0.715, 95% CI [−0.59, 0.41], d=−0.13, B_H(0,1)_=3.775; 90s, t(35)=−0.394, p=0.696, 95% CI [−0.60, 0.40], d=0.14, B_H(0,1)_=3.743; 120s, t(35)=−0.358, p=0.722, 95% CI [−0.59, 0.41], d=0.13, B_H(0,1)_=3.787. Therefore, there is substantial evidence for H_0_, with Bayes factors higher than 3, indicating that there are no differences in aSKNA between the two groups in the synchronous condition.

For the asynchronous condition, the results are as follows: 30s, t(35)=−0.365, p=0.717, 95% CI [−0.60, 0.42], d=0.12, B_H(0,1)_=3.779; 60s, t(35)=−0.381, p=0.705, 95% CI [−0.61, 0.41], d=0.13, B_H(0,1)_=3.759; 90s, t(35)=−0.393, p=0.697, 95% CI [−0.61, 0.42], d=0.14, B_H(0,1)_=3.745; 120s, t(35)=−0.381, p=0.705, 95% CI [−0.61, 0.42], d=0.13, B_H(0,1)_=3.759. Therefore, there is substantial evidence for H_0_, with Bayes factors higher than 3, indicating that there are no differences in aSKNA between the two groups in the asynchronous condition (Figure 4).

**Figure 4.**
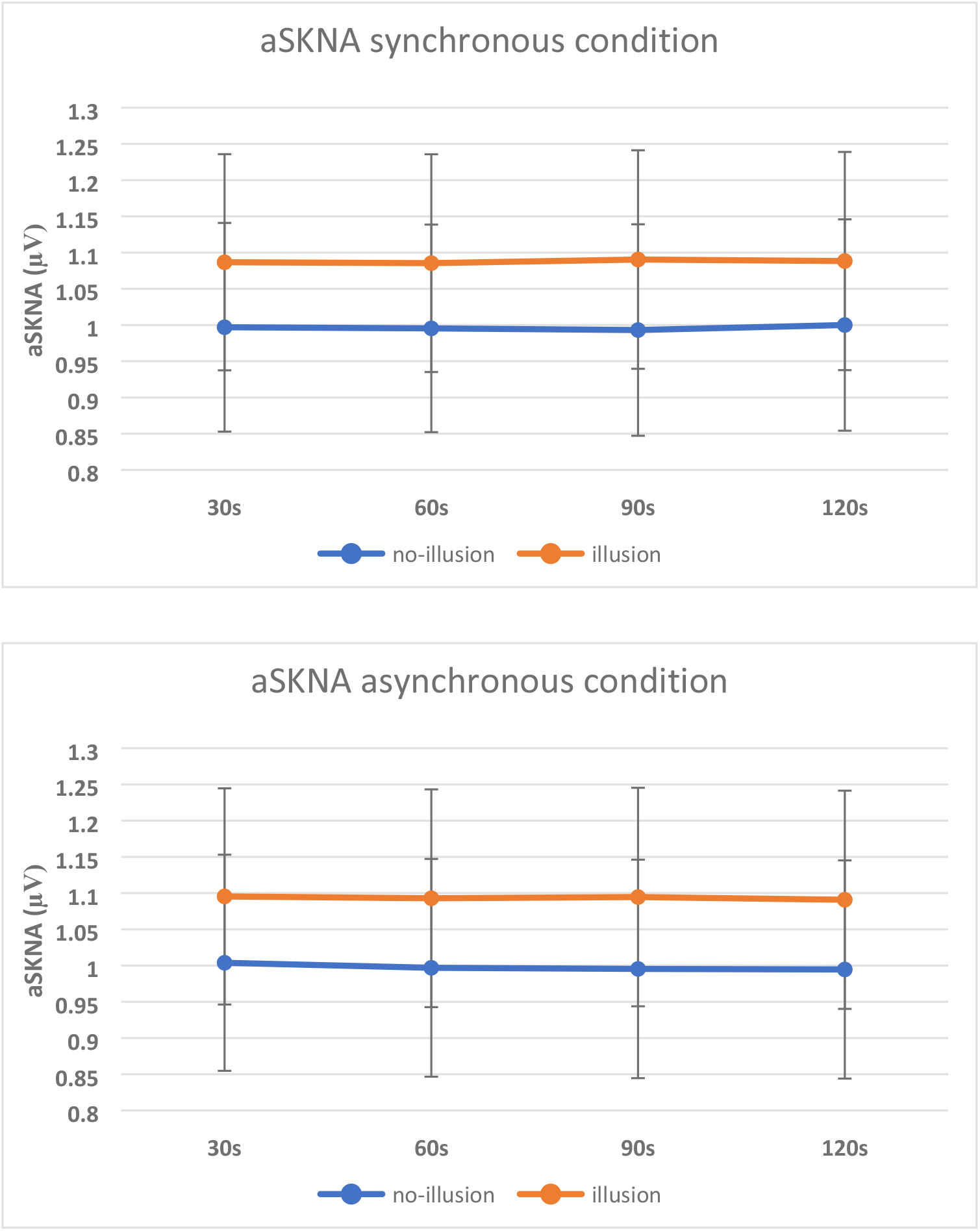
Average Skin nerve activity (aSKNA) in participants who were classified has having a strong versus minimal experience of the rubber hand illusion means ± 1SE. Following the methods of Doytchinova and colleagues [24], continuous recording of SNKA was processed to give average scores over the 30s for each synchronous (active – associated with rubber hand illusion) and asynchronous (control – not associated with the rubber hand illusion) stroking conditions of the task. No differences between individual who did and did not experience the illusion were observed for either the active or control condition.

#### Correlations across physiological measures

We computed correlations across autonomic measures, heart rate, heart rate variability, and skin conductance responses, and aSKNA for both synchronous and asynchronous conditions (Table 3). Overall these revealed little evidence for a systematic shift in the relationship between physiological variables that might arise form a different patterning of bodily control between synchronous and asynchronous conditions (implicitly linked to experience of the rubber hand illusion). We tested explicitly for the relationship between heart rate and heart rate variability. Typically, heart rate increases in heart rate are balanced by heart rate variability reflecting baroreflex activity. In stress/arousal states suppression of the baroreflex may change this relationship by inhibiting cardiovagal tone, decreasing heart rate variability and allowing heart rate and blood pressure to rise unchecked. We observed that during the asynchronous (control) condition heart rate was significantly negatively correlated with heart rate variability (R(35)=−0.593; p=0.000, 95CI [−0.769, −0.333] B=0.005). In the sychronous condition, associated with experience of the rubber hand illusion this relationship reduced in strength (R(35) −0.317, p = 0.056, 95CI [−0.581, 0.008] B=1.278). A test of the interaction, i.e. diffences in HR-HRV correlation between synchronous vs asynchronous conditions, did to reach significance Z=−1.460, p=0.144, indicating lack of compelling evidence across participants of a change in cardiac as a consequence of the rubber hand illusion induction (or effect).

**Table 3.**
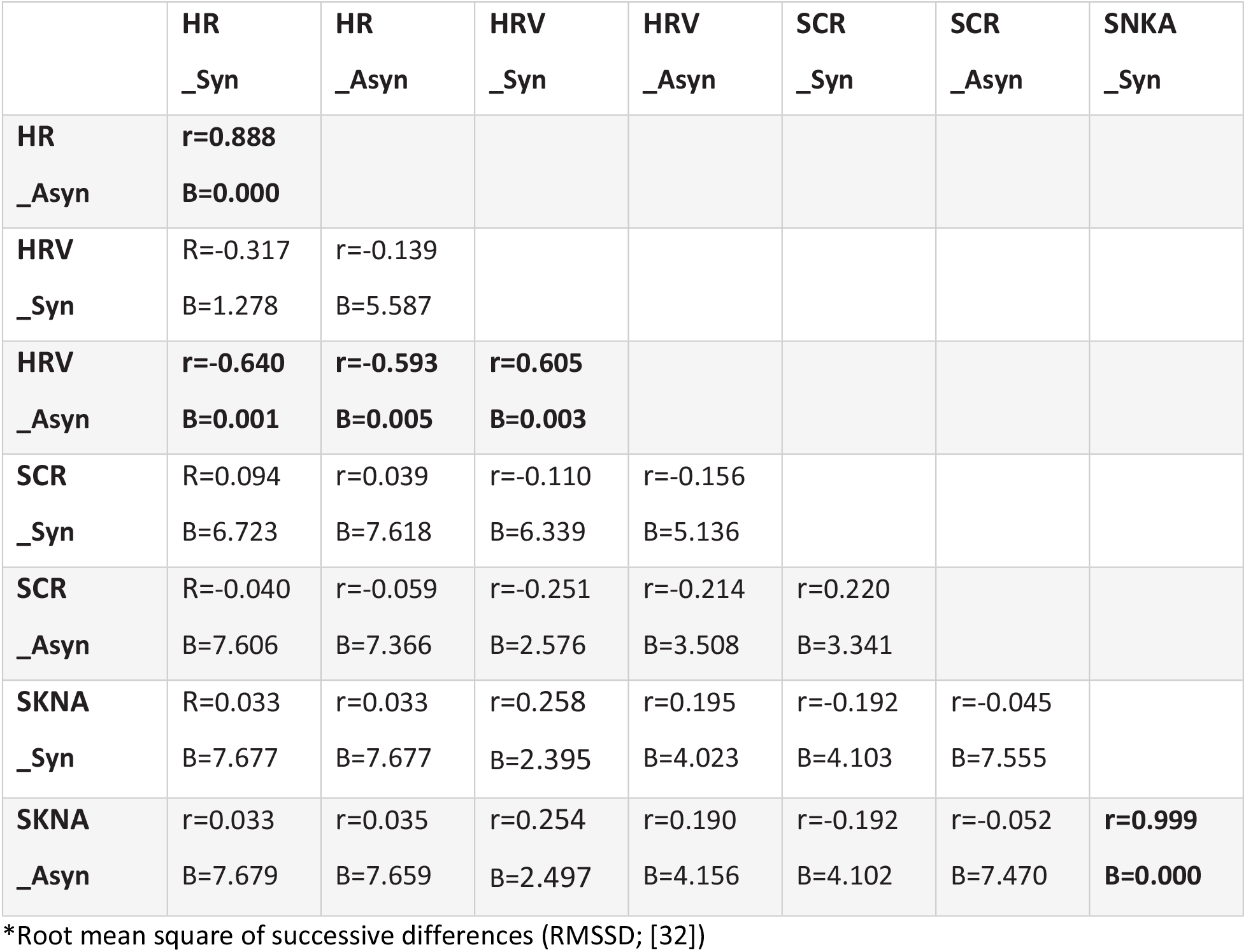
Correlations between Heart rate (HR), heart rate variability (HRV*), skin conductance responses (SCR) and aSKNA for synchronous (Syn) and asynchronous (Asyn) experimental conditions

|  | HR<br>_Syn | HR<br>_Asyn | HRV<br>_Syn | HRV<br>_Asyn | SCR<br>_Syn | SCR<br>_Asyn | SKNA<br>_Syn |
| --- | --- | --- | --- | --- | --- | --- | --- |
| HR<br>_Asyn | **r=0.888**<br>**B=0.000** |  |  |  |  |  |  |
| HRV<br>_Syn | R=-0.317<br>B=1.278 | r=-0.139<br>B=5.587 |  |  |  |  |  |
| HRV<br>_Asyn | **r=-0.640**<br>**B=0.001** | **r=-0.593**<br>**B=0.005** | **r=0.605**<br>**B=0.003** |  |  |  |  |
| SCR<br>_Syn | R=0.094<br>B=6.723 | r=0.039<br>B=7.618 | r=-0.110<br>B=6.339 | r=-0.156<br>B=5.136 |  |  |  |
| SCR<br>_Asyn | R=-0.040<br>B=7.606 | r=-0.059<br>B=7.366 | r=-0.251<br>B=2.576 | r=-0.214<br>B=3.508 | r=0.220<br>B=3.341 |  |  |
| SKNA<br>_Syn | R=0.033<br>B=7.677 | r=0.033<br>B=7.677 | r=0.258<br>B=2.395 | r=0.195<br>B=4.023 | r=-0.192<br>B=4.103 | r=-0.045<br>B=7.555 |  |
| SKNA<br>_Asyn | r=0.033<br>B=7.679 | r=0.035<br>B=7.659 | r=0.254<br>B=2.497 | r=0.190<br>B=4.156 | r=-0.192<br>B=4.102 | r=-0.052<br>B=7.470 | **r=0.999**<br>**B=0.000** |

\*Root mean square of successive differences (RMSSD; [32])

## Discussion

We set out to test whether the induction and experience of the rubber hand illusion is associated with reliable embodied changes in peripheral systemic (rather than localised) autonomic function linked theoretically to the notion of interoceptive predictive coding as a basis to the integrity of conscious self-representation. The initial ambition was to extract, from fine-grained recording of sympathetic nerves, a change in efferent autonomic nerve traffic that we hypothesised might encode a change in interoceptive prediction as a signature of the shift in the representation of the body and experience of self when an artificial limb is adopted as part of one’s own body. Our study incorporated two important features: first, the use of SKNA, a novel approach to record non-invasively from sympathetic nerves ([24, 26] and; second, the use of Bayes factors to determine strength of evidence for and against the null hypothesis [34]. In our findings, we produced robust evidence for successful induction of the rubber hand illusion, yet we observed no systematic changes across a number of autonomic measures during the induction procedure (synchronous stroking) relative to a control procedure (asynchronous stroking). Moreover, when participants were partitioned by a discriminant analysis as to whether or not the illusory experience was strong, autonomic measures in either condition did not distinguish between these groups. We also did not replicate a previously reported association between sensitivity to internal bodily signals (as indexed heuristically by performance accuracy on a heartbeat tracking task) and resistance to the rubber hand illusion.

Retrospectively, seeing a bodily signature of the rubber hand illusion was perhaps ambitious. We pursued autonomic measures that are expressed thought the body (electrodermal, cardiovascular and SKNA), rather than more limb-specific (previous reports of hand cooling associated with rubber hand illusion found this to be localised to the specific side of the illusion). Our findings of no sympathetic electrodermal or SKNA changes associated with the rubber hand illusion contrast with recent observations that differences in skin conductance occur during the rubber hand illusion [19]. In this study, transient effects were observed, suggesting the novelty of the experience was a contributing factor. Increases in phasic and tonic electrodermal activity were also reported in an earlier study, here associated with the period leading up to the onset of the illusory experience of body ownership, and amplified in people who report high levels of anomalous bodily experiences [35]. Both these studies suggest that autonomic changes, if they occur, may be brief, and mostly related to the cognitive and emotional experience of the induction process.

The rubber hand illusion studies that report a fall in skin temperature in the ‘replaced hand’ have been influential and widely cited as evidence for a deep physical embodiment [15]. Temperature after a 7–8 min stroking period is reported as lower with synchronous compared to asynchronous stroking on the test hand. However, whereas no temperature difference is reported on the non-stimulated hand. This observation suggests that hand cooling might relate to the illusionary disowning of the real hand in favour of the rubber substitute. Alternative, yet weaker explanations include greater immobility of the target hand (during synchronous stroking) and/or the illusion) features of the synchronous stoking that might better elicit peripheral e.g. axo-axonal reflex hypo-perfusion. Social factors may also play a role [18]. Mechanisms for the observed hand cooling are relevant to other observations: While passive disowning of the replaced hand most likely would be accompanied by reduced autonomic innervation of the limb, the increase in electrodermal activity occasionally reported with the rubber hand illusion is also consistent with activation of sympathetic vasoconstrictor neurons within skin vasculature which would also actively reduce skin temperature [36]. A number of studies fail to find a cooling effect associated with the rubber hand illusion (e.g.[17]). In our study, there was strong evidence for no effect of the procedure or the illusion on electrodermal activity or skin sympathetic activity measured from the chest surface. Nevertheless, our induction process was relatively short; it is possible that transient changes at the onset of the illusion were not sustained long enough to affect the average skin conductance data. For SKNA measures, we did analyse these sensitive response data for shorter time periods over the course of the experiment and yet still identified no effect to suggest a transient response.

We measured heart rate and heart rate variability (RMSSD) also, both indices of the autonomic cardiac control capturing the interaction of sympathetic and parasympathetic (vagus nerve) cardiac innervation [31, 32]. Heart rate variability in particularly is a widely used measure linked to health, well-being and cognitive flexibility, where decreases in heart rate variability accompany stress, anxiety and negative affective experience [37, 38]. These decreases relate to a suppression of the baroreflex by top-down brain signals, enabling heart rate and blood pressure to rise together to meet behavioural allostatic demands. Yet again, within our data, we did not find any clear association with the experience of illusory limb ownership. Heart rate itself is often used as a rough proxy for sympathetic cardiac influences; heart rate typically increases with decreasing heart rate variability as changes in sympathovagal balance produce cardiovascular arousal.

Attempts to quantify sympathetic nerve traffic more directly include the established technique of microneurography [23, 39–41], which employs microelectrodes inserted into peripheral nerves (median or common peroneal) to measure multiunit activity (sometimes also single unit action potentials) from sympathetic axon bundles. Muscle sympathetic nerve activity shows different patterns of reactivity compared to skin sympathetic, reflecting the dependence on the baroreflex in gating sympathetic vascular control [23]. The relatively recent introduction of non-invasive skin sympathetic nerve activity (SKNA) recordings provides a means of overcoming some of the logistical limitations of invasive microneurography [24–26, 42]. Subcutaneous nerve activity was initially detected with implanted bipolar electrodes in dogs and, when recorded from the chest wall, was found to correlated with the stellate ganglion nerve activity and sympathetic nervous effects on heart [43]. SKNA (differentiated from the skin sympathetic nerve activity underpinning electrodermal responses) is reportedly a more sensitive than heart rate variability derived indices of cardiac sympathetic tone [44]. It was subsequently shown that this could be recorded in humans with standard electrocardiographic electrode and high sample rates. Validation studies in humans have established the sensitivity of SKNA recordings to autonomic challenges in healthy individuals, to lidocaine deactivation of stellate ganglion, to vagus nerve stimulation, and to pathological changes in cardiac rhythm [24, 26, 42]. As a measure of multiunit sympathetic activity relevant to systemic cardiovascular control, this approach is exciting for both cardiological investigations and for human behavioural psychophysiology. This is the first instantiation of this approach in such an experimental context. We followed published methods for recording SKNA [24, 26], setting out to leverage this promising technique to provide higher quality data on efferent neural signatures of central changes in self-representation. However, we report no differences in this measure between the active (synchronous stroking) and control (asynchronous stroking) conditions of the rubber hand illusion no whether a strong experience of change in embodied representation occurred.

One interesting challenge within the field of body-brain Interaction (including illusions of body ownership, interoception and autonomic psychophysiology) is the pursuit of objective measures, unbiased by situational and individual confounds such implicit task demands, response bias, and suggestibility. Tests of interoceptive sensitivity, especially the heartbeat tracking task have been criticised from the perspective of psychophysics for not providing an unconfounded measure of sensory perception; individual difference is performance are subject to a number of non-sensory factors including knowledge of one’s heart rate practice effects and other top down beliefs. These widely acknowledged associations are of lesser or greater relevance depending on what question the research study aims to address and correspondingly, whether the data are interpreted appropriately. Often what is sought is a predictor of a psychological trait symptom or vulnerability such a performance biases in subjective representation/appraisal of bodily signals may be more proximate to the target experience. Relevant to this study, individual differences in social compliance, suggestibility and /or neuroticism are long-recognized factors determinants of behaviours relating to ‘strength of self-representation’. Blinding / double-blinding of clinic trials represent an attempt to manage unwanted influences of expectancies and biases that permeate research questionnaires and the structure of experiments. Suggestibility, including hypnotizability, and phenomenological control influence measures of ‘objective’ performance on tasks designed to access conscious processes, including synaesthetic perceptual experience, interoceptive ability, free-will action generation, and the subjective experience of the rubber hand illusion. Individual differences in suggestibility can may account for around 10 percent of the variance in subjective measures of the rubber hand illusion [45]. Such observations indicate that superordinate domain-general beliefs and expectancies unsurprisingly influence perceptual behavioural and experiential measures. Moreover, they highlight the importance of participant-level individual lability and experiment-level contextual coerciveness in shaping subjective experience. In this study we did not assess if suggestibility influenced autonomic responses to induction or experience of the rubber had illusion, beyond using the established standard asynchronous stroking as a control condition. Our autonomic data showing a lack of difference between synchronous and asynchronous condition or participants that are like to insensitive to address the question whether suggestion effects autonomic signals (as can be evoked with visual imagery [46]). Moreover, we did not see effects associated with individual differences in performance of a heartbeat counting task, a heuristic measure of perceptual sensitivity to internal bodily cues that has been criticised for its sensitivity to task demands and related top-down or contextual effects. Increased performance on this task has been linked to enhanced autonomic reactivity [47] and to resistance to rubber hand illusion [12]. A previous study also did not replicate this latter effect showing a stronger associate with affective touch [48].

In conclusion, we undertook detailed autonomic monitoring of participants during the rubber hand illusion and found no systematic effect on sympathetic or cardiovagal measures that might discriminate between the induction procedure or state of ownership of an illusory body part versus a control condition. We obtained, for the first time in this context, SKNA recordings, indexing skin sympathetic nerve activity proximally related to stellate ganglion activity. We used Bayes factors to confirm the absence of autonomic differences. Our study illustrates that tonic changes in bodily physiology and not obligatory accompaniments of an alteration in his type of conscious self-representation.

## Acknowledgements

This study was funded by a bursary from the Bial Foundation 128/14 to HC entitled: *Microneurography and autonomic nerve recording as a tool for consciousness science.* We are grateful for the advice and help of Dr David Watson in setting up the data acquisition platform, and to Prof Peter Taggart for highlighting and advising on the SKNA methodology.

## References

1 Blanke O. Multisensory brain mechanisms of bodily self-consciousness. Nature Reviews Neuroscience 2012, 13, 556–571.

2 Gallagher S. Philosophical conceptions of the self: Implications for cognitive science. Trends in Cognitive Sciences 2000. 4, 14–21.

3 van den Bos E & Jeannerod M. Sense of body and sense of action both contribute to self-recognition. Cognition2002, 85, 177–187.

4 Botvinick M & Cohen J Rubber hands ‘feel’ touch that eyes see. Nature 1998, 391, 756.

5 Armel KC & Ramachandran VS. Projecting sensations to external objects: Evidence from skin conductance response. Proceedings of the Royal Society of London, Series B: Biological Sciences 2003, 270, 1499–1506.

6 Kilteni K, Normand J-M, Sanchez-Vives MV, Slater M. Extending body space in immersive virtual reality: A very long arm illusion. PLoS One 2012, 7, e40867.

7 Damasio AR. The feeling of what happens: Body and emotion in the making of consciousness. Houghton Mifflin Harcourt. 1999.

8 Craig AD. How do you feel—now? The anterior insula and human awareness. Nature Reviews Neuroscience 2009, 10, 59–70.

9 Seth A, Tsakiris M Being a Beast Machine: The Somatic Basis of Selfhood Trends in Cognitive Sciences 2018 22: 969–981.

10 Sierra M, Senior C, Dalton J, McDonough M, Bond A, Phillips ML, O’Dwyer AM, David AS. Autonomic response in depersonalization disorder. Arch Gen Psychiatry. 2002 59(9):833–8. doi: 10.1001/archpsyc.59.9.833.

11 Koreki A, Garfkinel SN, Mula M, Agrawal N, Cope S, Eilon T, Gould Van Praag C, Critchley HD, Edwards M, Yogarajah M. Trait and state interoceptive abnormalities are associated with dissociation and seizure frequency in patients with functional seizures. Epilepsia 2020. 61:1156–1165. doi: 10.1111/epi.16532.

12 Tsakiris M, Tajadura-Jiménez A, & Costantini M. Just a heartbeat away from one’s body: Interoceptive sensitivity predicts malleability of body-representations. Proceedings of the Royal Society of London, Series B: Biological Sciences 2011, 278, 2470–2476.

13 Suzuki K, Garfinkel SN, Critchley HD & Seth AK. Multisensory integration across exteroceptive and interoceptive domains modulates self-experience in the rubber-hand illusion. Neuropsychologia 2013, 51, 2909–2917.

14 Aspell JE, Heydrich L, Marillier G, Lavanchy T, Herbelin B, Blanke O. Turning body and self inside out: visualized heartbeats alter bodily self-consciousness and tactile perception. Psychol Sci 2013. 24:2445–53.

15 Moseley GL, Olthof N, Venema A, Don S, Wijers M, Gallace A, Spence C. Psychologically induced cooling of a specific body part caused by the illusory ownership of an artificial counterpart. Proceedings of the National Academy of Sciences, U.S.A 2008 105, 13169–13173.

16 Kammers MPM, Rose K, Haggard P. Feeling numb: temperature, but not thermal pain, modulates feeling of body ownership. Neuropsychologia 2011. 49: 1316–1321.

17 de Haan AM, Van Stralen HE, Smit M, Keizer A, Van der Stigchel S, & Dijkerman HC. No consistent cooling of the real hand in the rubber hand illusion. Acta Psychologica 2017, 179, 68–77.

18 Rohde M, Wold A, Karnath HO, Ernst MO. The human touch: skin temperature during the rubber hand illusion in manual and automated stroking procedures. PLoS One2013. 8(11):e80688.

19 D’Alonzo M, Mioli A, Formica D, Di Pino G. Modulation of Body Representation Impacts on Efferent Autonomic Activity. J Cogn Neurosci. 2020 32(6):1104–1116.

20 Braithwaite J, Watson DG, Jones R, Rowe M. A guide for analysing electrodermal activity (EDA) & skin conductance responses (SCRs) for psychological experiments. Psychophysiology 2013, 49(1), 1017–1034.

21 Boucsein, W. Electrodermal activity (2nd ed.). Springer Science + Business Media. 2012 https://doi.org/10.1007/978-1-4614-1126-0.

22 Critchley HD, Elliott R, Mathias CJ, Dolan RJ. Neural activity relating to generation and representation of galvanic skin conductance responses: a functional magnetic resonance imaging study. J Neurosci 2000. 20: 3033–40.

23 Wallin BG, Donadio V, Karlsson T, Kallio M, Nordin M, Elam M. Arousal increases baroreflex inhibition of muscle sympathetic activity. Acta Physiol Scand 2003. 177(3):291–8.

24 Doytchinova A, Hassel JL, Yuan Y, Lin H, Yin D, Adams D, Straka S, Wright K, Smith K, Wagner D, Shen C, Salanova V, Meshberger C, Chen LS, Kincaid JC, Coffey AC, Wu G, Li Y, Kovacs RJ, Everett TH 4th, Victor R, Cha YM, Lin SF, Chen PS. Simultaneous noninvasive recording of skin sympathetic nerve activity and electrocardiogram. Heart Rhythm 2017. 14(1):25–33.

25 Everett TH 4th, Doytchinova A, Cha YM, Chen PS. Recording sympathetic nerve activity from the skin. Trends Cardiovasc Med 2017. 27(7):463–472.

26 Kusayama T, Wong J, Liu X, He W, Doytchinova A, Robinson EA, Adams DE, Chen LS, Lin SF, Davoren K, Victor RG, Cai C, Dai MY, Tian Y, Zhang P, Ernst D, Rho RH, Chen M, Cha YM, Walega DR, Everett TH 4th, Chen PS. Simultaneous noninvasive recording of electrocardiogram and skin sympathetic nerve activity (neuECG). Nat Protoc 2020. 15(5):1853–1877.

27 Longo MR, Schüür F, Kammers MP, Tsakiris M & Haggard P. What is embodiment? A psychometric approach. Cognition 2008 107(3), 978–998.

28 Schandry,R. Heart beat perception and emotional experience. Psychophysiology 1981, 18(4), 483–488.

29 Garfinkel SN, Seth AK, Barrett AB, Suzuki K, Critchley HD. Knowing your own heart: distinguishing interoceptive accuracy from interoceptive awareness. Biological Psychology 2015, 104, 65–74.

30 Fedotov AA. Selection of parameters of bandpass filtering of the ECG signal for heart rhythm monitoring systems. Biomedical Engineering 2016, 50(2), 114–118.

31 Shaffer F & Ginsberg JP. An overview of heart rate variability metrics and norms. Frontiers in Public Health 2017, 5, 258.

32 Munoz ML, van Roon A, Riese H, Thio C, Oostenbroek E, Westrik I, de Geus EJ, Gansevoort R, Lefrandt J, Nolte IM, Snieder H. (Validity of (Ultra-)Short Recordings for Heart Rate Variability Measurements. PLoS One2015. 10(9):e0138921

33 Garfinkel SN, Minati L, Gray MA, Seth AK, Dolan RJ, & Critchley HD. Fear from the heart: sensitivity to fear stimuli depends on individual heartbeats. Journal of Neuroscience 2014, 34(19), 6573–6582.

34 Keysers C, Gazzola V, Wagenmakers EJ. Using Bayes factor hypothesis testing in neuroscience to establish evidence of absence. Nat Neurosci. 202023(7):788–799.

35 Braithwaite JJ, Broglia E, Watson DG. Autonomic emotional responses to the induction of the rubber-hand illusion in those that report anomalous bodily experiences: Evidence for specific psychophysiological components associated with illusory body representations · J Exp Psychol Hum Percept Perform 2014. 40(3): 1131–45.

36 Bini G, Hagbarth KE, Hynninen P, Wallin BG Thermoregulatory and rhythm-generating mechanisms governing the sudomotor and vasoconstrictor outflow in human cutaneous nerves J Physiol (Lond) 1980, 306, 537–552.

37 Drury RL, Porges S, Thayer J, Ginsberg JP. Heart Rate Variability, Health and Well-Being: A Systems Perspective. Front Public Health 2019. 7:323.

38 Mulcahy JS, Larsson DEO, Garfinkel SN, Critchley HD. Heart rate variability as a biomarker in health and affective disorders: A perspective on neuroimaging studies. Neuroimage 2019. 202:116072

39 Sverrisdóttir YB, Schultz T, Omerovic E, Elam M. Sympathetic nerve activity in stress-induced cardiomyopathy. Clin Auton Res 2012. 22(6):259–64.

40 Vallbo ÅB. Microneurography: how it started and how it works. J Neurophysiol. 2018 120(3):1415–1427.

41 Macefield VG. Recording and quantifying sympathetic outflow to muscle and skin in humans: methods, caveats and challenges. Clin Auton Res. 2020 doi: 10.1007/s10286-020-00700-6.

42 Kusayama T, Wan J, Doytchinova A, Wong J, Kabir RA, Mitscher G, Straka S, Shen C, Everett TH 4th, Chen PS. Skin sympathetic nerve activity and the temporal clustering of cardiac arrhythmias. JCI Insight 2019. 4(4):e125853.

43 Jiang Z, Zhao Y, Doytchinova A, Kamp NJ, Tsai WC, Yuan Y, Adams D, Wagner D, Shen C, Chen LS, Everett TH 4th, Lin SF, Chen PS. (Using skin sympathetic nerve activity to estimate stellate ganglion nerve activity in dogs. Heart Rhythm 2015. 12(6):1324–32.

44 Chan YH, Tsai WC, Shen C, Han S, Chen LS, Lin SF, Chen PS. Subcutaneous nerve activity is more accurate than heart rate variability in estimating cardiac sympathetic tone in ambulatory dogs with myocardial infarction. Heart Rhythm 2015. 12(7): 1619–27.

45 Lush P, Botan V, Scott RB, Ward J, Dienes Z. Phenomenological control: response to imaginative suggestion predicts measures of mirror touch synaesthesia, vicarious pain and the rubber hand illusion Nature Comms 2020 in press [2019. psyarxiv.com].

46 Wang Y, Morgan WP. The effect of imagery perspectives on the psychophysiological responses to imagined exercise. Behav Brain Res. 1992 52(2):167–74

47 Pollatos O, Herbert BM, Kaufmann C, Auer DP, Schandry R. Interoceptive awareness, anxiety and cardiovascular reactivity to isometric exercise. Int J Psychophysiol 2007. 65(2):167–73.

48 Crucianelli L, Krahé C, Jenkinson PM, Fotopoulou AK. Interoceptive ingredients of body ownership: Affective touch. Cortex 2018 104:180–192.

